## Supplemental Figures 1-10 for "The role of satellite cell-derived TRIM28 in mechanical load- and injury-induced myogenesis"

Kuan-Hung Lin *et al.*

**This PDF file includes:**

Figs. S1 to S10
Tables S1

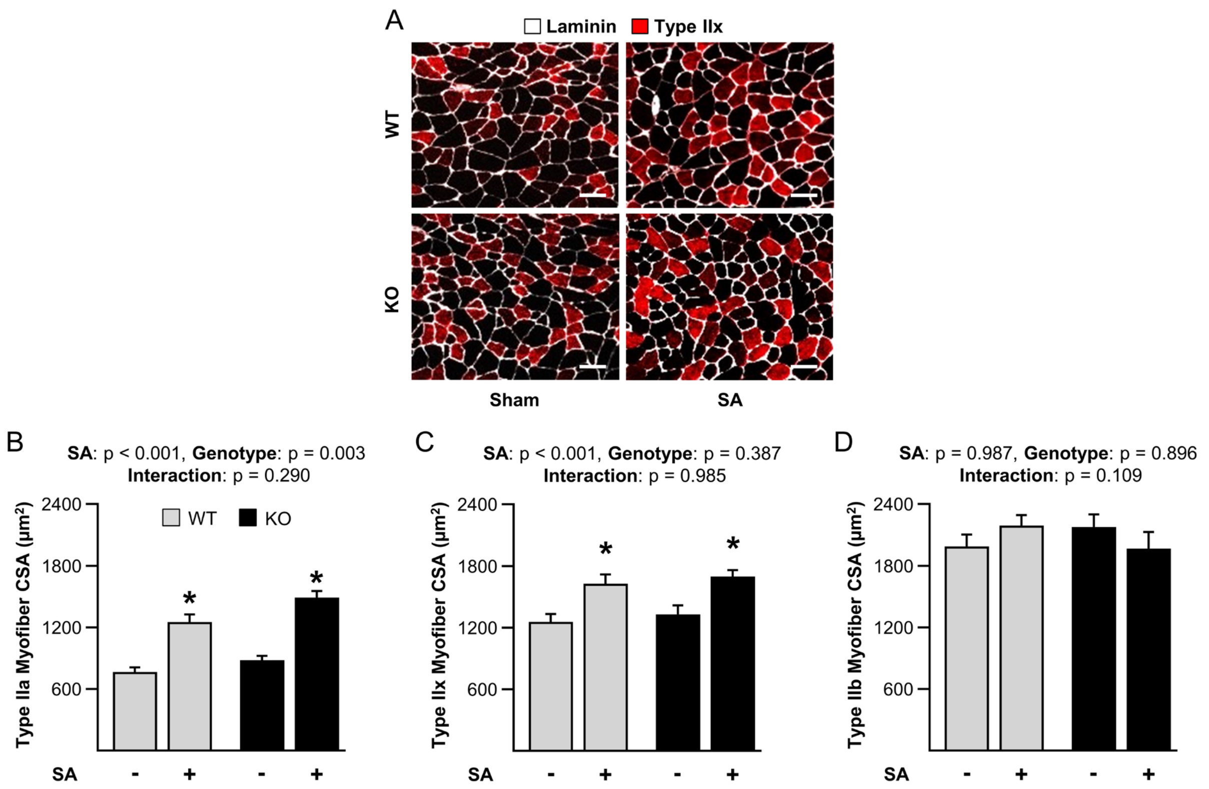


**Fig. S1. The loss of TRIM28 in satellite cells does not impact the mechanical load-induced increase in the size of Type IIa or IIx myofibers.** Wild-type (WT) mice and tamoxifen-inducible satellite cell-specific TRIM28 knockout mice (KO) mice were treated with tamoxifen. At 14 days post tamoxifen, mice were subjected to unilateral synergist ablation surgery (SA+), with the non-ablated limb serving as a sham control (SA-). The mice were treated as described in Fig. 3 and the plantaris muscles were collected at 14 days after the SA surgery. (**A**) Mid-belly cross-sections were subjected to immunohistochemistry for laminin and type IIx myofibers. (**B**-**D**) Quantification of the type IIa, type IIx, and type IIb myofiber cross-sectional area, respectively. Values are group means + SEM, *n* = 11-12/group. Data were analyzed with two-way ANOVA. * indicates a significant effect of SA within the given genotype, $p < 0.05$. Scale bars = 50 µm.

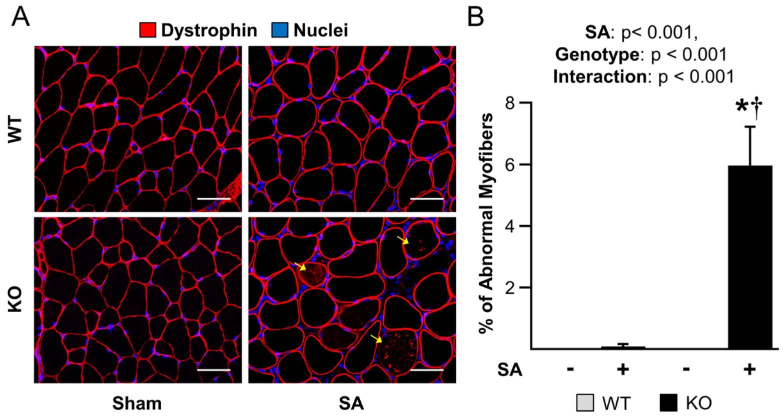


**Fig. S2. The loss of TRIM28 in satellite cells leads to the presence of myofibers with abnormal dystrophin at 14 days after the onset of synergist ablation.** Wild-type (WT) mice and tamoxifen-inducible satellite cell-specific TRIM28 knockout mice (KO) mice were treated with tamoxifen. At 14 days post tamoxifen, mice were subjected to unilateral synergist ablation surgery (SA+), with the non-ablated limb serving as a sham control (SA-). The mice were treated as described in Fig. 3 and the plantaris muscles were collected at 14 days after the SA surgery. (**A**) Mid-belly cross-sections were subjected to immunohistochemistry for dystrophin and nuclei. Yellow arrows indicate dystrophin detected within myofibers. (**B**) Quantification of myofibers that had an abnormal appearance of dystrophin. Values are group means + SEM, *n* = 6/group. Data were analyzed with two-way ANOVA. * indicates a significant effect of SA within the given genotype, † indicates a significant difference between the SA groups, $p < 0.05$. Scale bars = 50 µm.

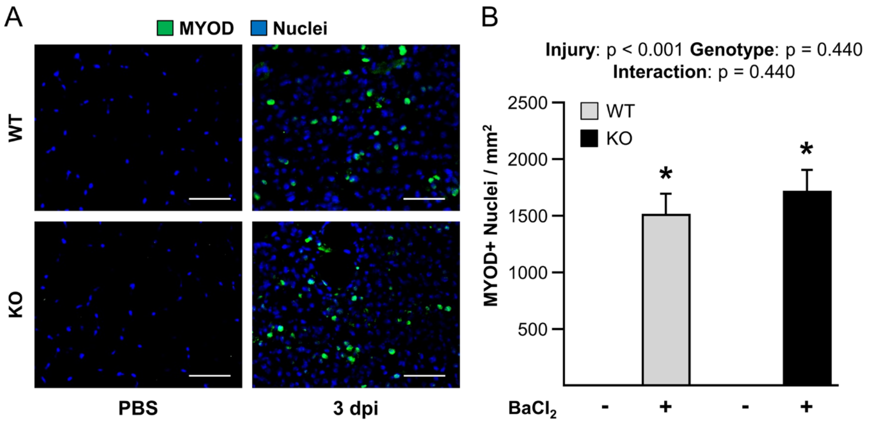


**Fig. S3. The loss of TRIM28 in satellite cells does not impair injury-induced satellite cell activation.** Wild-type (WT) mice and tamoxifen-inducible satellite cell-specific TRIM28 knockout mice (KO) mice were treated with tamoxifen. At 14 days post tamoxifen, their tibialis anterior muscles were injected with $BaCl_2$ (+) to induce injury or PBS (-) as a control condition. (**A**) The tibialis anterior muscles were collected at 3 days post-injury (dpi) and then mid-belly cross-sections were subjected to immunohistochemistry for MYOD (a marker of satellite cell activation), and nuclei. (**B**) Quantification of the number of MYOD-positive nuclei per $mm^2$ in A. Values are group means + SEM, *n* = 4/group. Data were analyzed with two-way ANOVA. * indicates a significant effect of injury within a given genotype, † indicates a significant difference between the $BaCl_2$ treated groups, $p < 0.05$. Scale bars = 50 μm.

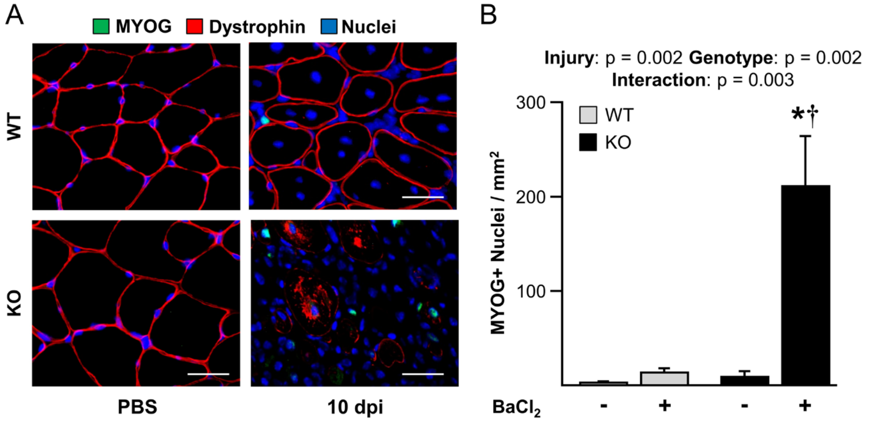


**Fig. S4. The loss of TRIM28 in satellite cells does not impair injury-induced satellite cell differentiation.** Wild-type (WT) mice and tamoxifen-inducible satellite cell-specific TRIM28 knockout mice (KO) mice were treated with tamoxifen. At 14 days post tamoxifen, their tibialis anterior muscles were injected with $BaCl_2$ (+) to induce injury or PBS (-) as a control condition. (**A**) The tibialis anterior muscles were collected at 10 days post-injury (dpi) and then mid-belly cross-sections were subjected to immunohistochemistry for MYOG, dystrophin, and nuclei. (**B**) Quantification of the number of MYOG-positive nuclei per $mm^2$ in A. Values are group means + SEM, $n$ = 4/group. Data were analyzed with two-way ANOVA. * indicates a significant effect of injury within a given genotype, † indicates a significant difference between the $BaCl_2$ treated groups, $p < 0.05$. Scale bars = 50 μm.

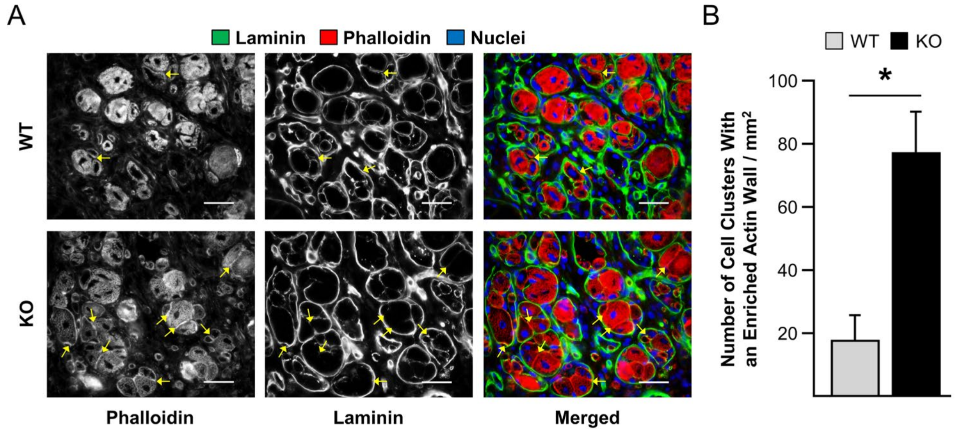


**Fig. S5. The loss of TRIM28 leads to an increase in the presence of dense walls of filamentous actin between well-aligned myoblasts/myofibers.** Wild-type (WT) mice and tamoxifen-inducible satellite cell-specific TRIM28 knockout mice (KO) mice were treated with tamoxifen. At 14 days post tamoxifen, their tibialis anterior (TA) muscles were injected with $BaCl_2$ to induce injury. (**A**) The tibialis anterior muscles were collected at 7 days post-injury and mid-belly cross-sections were subjected to immunohistochemistry for laminin, filamentous actin (i.e., phalloidin), and nuclei. (**B**) Clusters of well-aligned phalloidin-positive myoblasts/myofibers that were surrounded by a thick outer layer of laminin were identified, and then the number of clusters that contained phalloidin-positive myoblasts/myofibers with a dense wall of filamentous actin between the aligned cells (yellow arrows) was quantified. Values are group means + SEM, $n$ = 4/group. Data were analyzed with a Student's t-test. * indicates a significant difference between groups. $p < 0.05$. Scale bars = 50 µm.

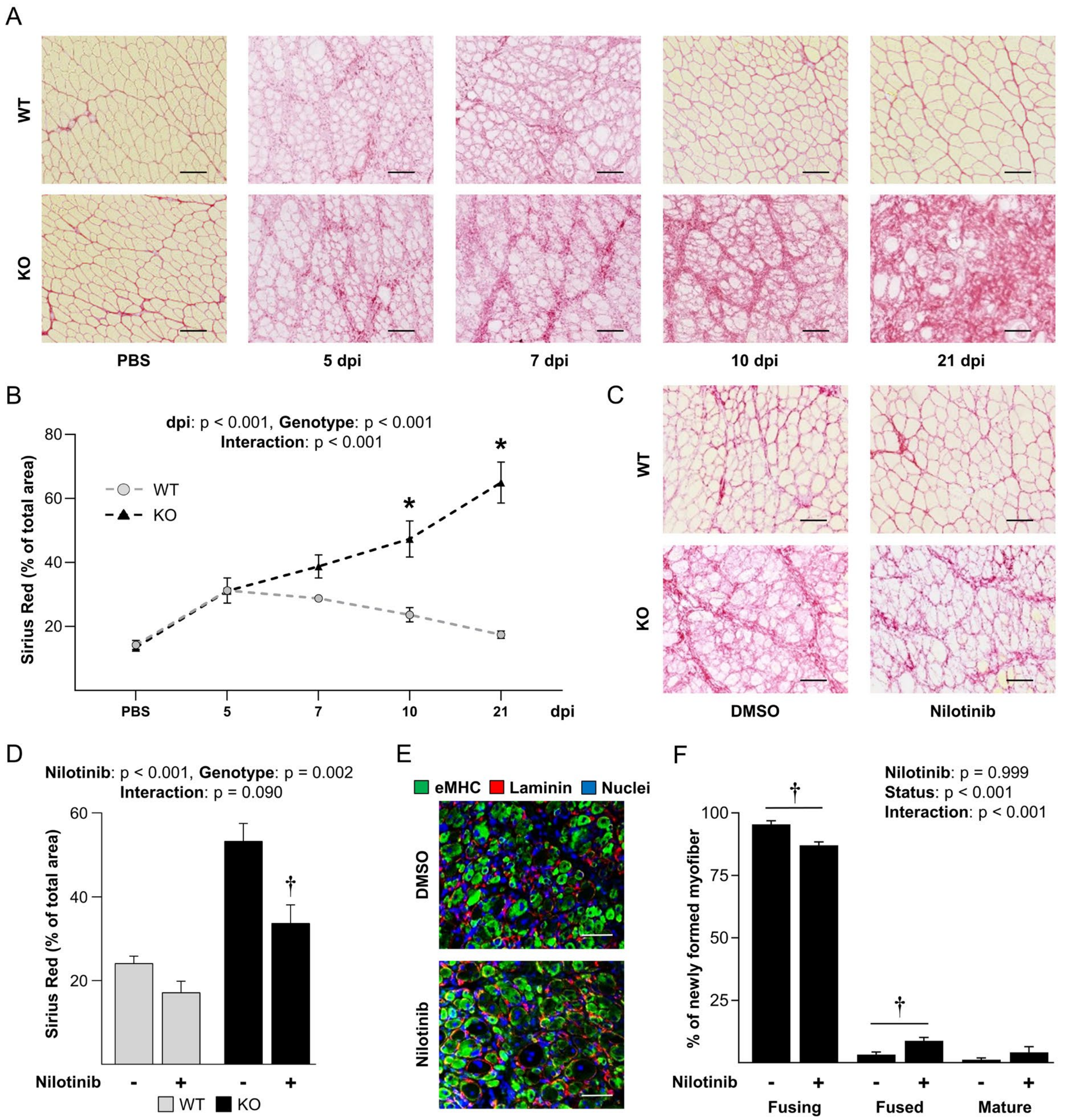


**Fig. S6. The loss of TRIM28 in satellite cells leads to excessive fibrosis following $BaCl_2$-induced injury, but reducing the fibrosis only minimally improves the impairment in fusion.** Wild-type (WT) mice and tamoxifen-inducible satellite cell-specific TRIM28 knockout mice (KO) mice were treated with tamoxifen. At 14 days post tamoxifen, their tibialis anterior (TA) muscles were injected with $BaCl_2$ to induce injury or PBS as a control condition. (**A**) At 5, 7, 10, and 21 days post-injury (dpi), TA muscles were collected and mid-belly cross-sections were subjected to Sirius Red staining as a marker of collagen deposition. (**B**) Proportion of the muscle cross-section that stained positive for Sirius Red in A. (**C-F**) Daily intraperitoneal injections of Nilotinib (20 mg/kg/day) or DMSO were administered at 3 to 7 dpi. (C) TA muscles were collected at 10 dpi and mid-belly cross-sections were subjected to Sirius Red staining. (D) Proportion of the muscle cross-sections that stained positive for Sirius Red in C. (E) Mid-belly cross-sections of the TA muscles at 10 dpi were subjected to immunohistochemistry for eMHC, laminin, and nuclei. (F)

Proportion of myoblasts/myofibers that were at the stage of fusing, fused, or mature as described in Fig. 4. Values are group means + SEM, $n$ = 4-6/group. Data were analyzed with two-way ANOVA. * indicates a significant difference between genotypes at the given timepoint, † indicates a significant effect of Nilotinib within the given genotype or stage, $p < 0.05$. Scale bars = 100 μm in A and C, and 50 μm in E.

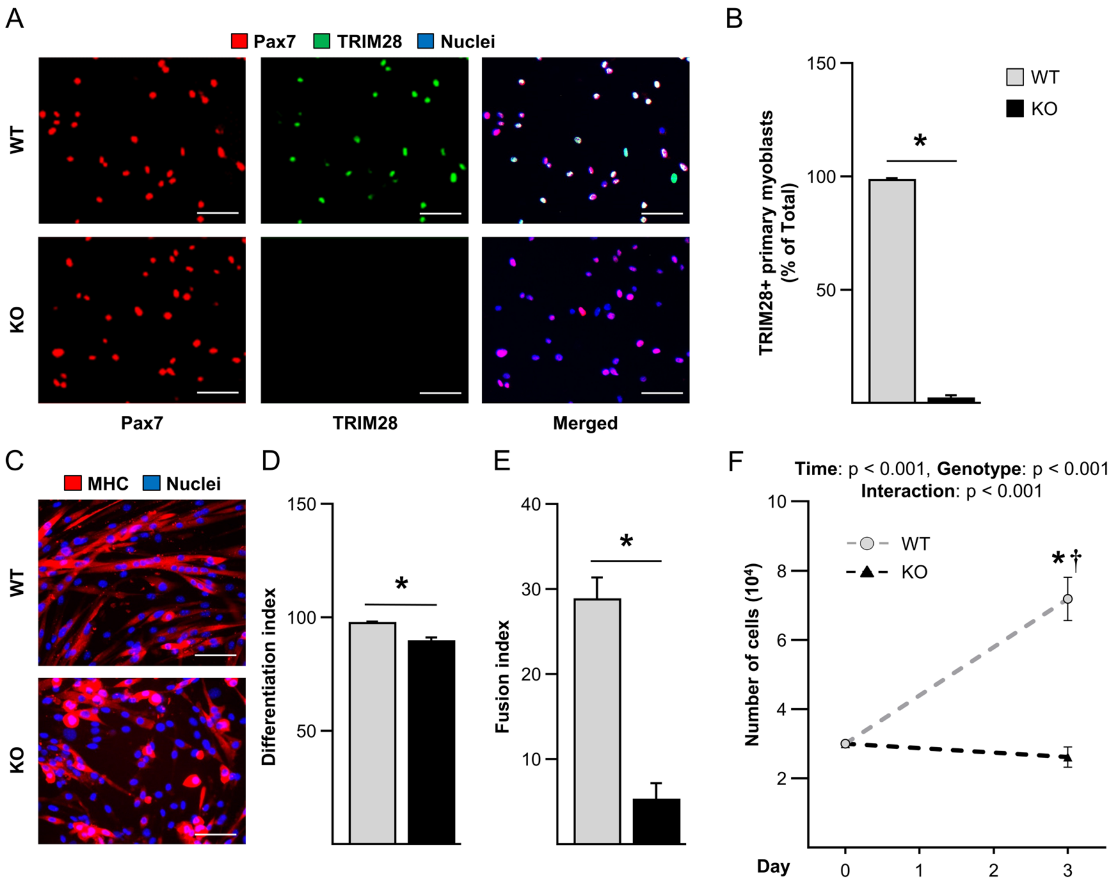


**Fig. S7. The loss of TRIM28 in satellite cells leads to a fusion defect during *in vitro* myotube formation.** Wild-type (WT) mice and tamoxifen-inducible satellite cell-specific TRIM28 knockout mice (KO) mice were treated with tamoxifen. (**A**) At 14 days post tamoxifen, primary myoblasts were isolated, cultured in growth medium, and then subjected to immunohistochemistry for Pax7, TRIM28, and nuclei. (**B**) The proportion of the Pax7 positive primary myoblasts that expressed TRIM28 in A. (**C-E**) WT and KO primary myoblasts were subjected to a myotube formation assay and immunohistochemistry for myosin heavy chain (MHC) and nuclei. (D) The differentiation index (% of nuclei inside MHC positive cells), and (E) the fusion index (% of nuclei inside MHC positive multinucleated cells) were quantified. (**F**) $3x10^4$ primary myoblasts were seeded on day 0 and cultured in growth medium. The number of primary myoblasts was quantified on day 3. Values are group means + SEM, *n* = 4-6/group with each sample representing an independent line of isolated primary myoblasts. Data in B, D, and E were analyzed with a Student's t-test. Data in F were analyzed with two-way repeated-measures ANOVA. * indicates a significant difference between genotypes within a given condition, † indicates a significant difference from day 0, $p < 0.05$, Scale bars = 50 μm.

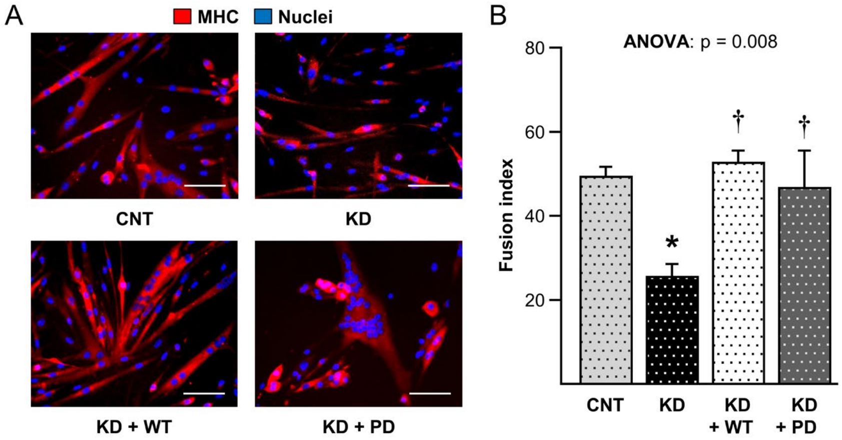


**Fig. S8. TRIM28(S473) phosphorylation is not required for fusion during *in vitro* myotube formation.** Primary myoblasts were infected with lentivirus encoding scrambled shRNA (CNT), shRNA targeting *Trim28* mRNA (KD), shRNA targeting *Trim28* mRNA along with "rescue" expression of a shRNA resistant form of TRIM28 (KD + WT), or shRNA targeting *Trim28* mRNA along with "rescue" expression of an shRNA resistant and phosphodefective form of TRIM28 (KD + PD) in which the serine 473 residue had been mutated to a non-phosphorylatable alanine. (**A**) Infected myoblasts were subjected to a myotube formation assay and immunohistochemistry for MHC and nuclei. (**B**) Quantification of the fusion index (% of nuclei inside MHC positive and multinucleated cells). Values are group means + SEM, $n$ = 4/group with each sample representing an independent line of isolated primary myoblasts. Data were analyzed with one-way ANOVA. * indicates a significant difference from CNT, † indicates a significant difference from KD, $p < 0.05$. Scale bars = 50 μm.

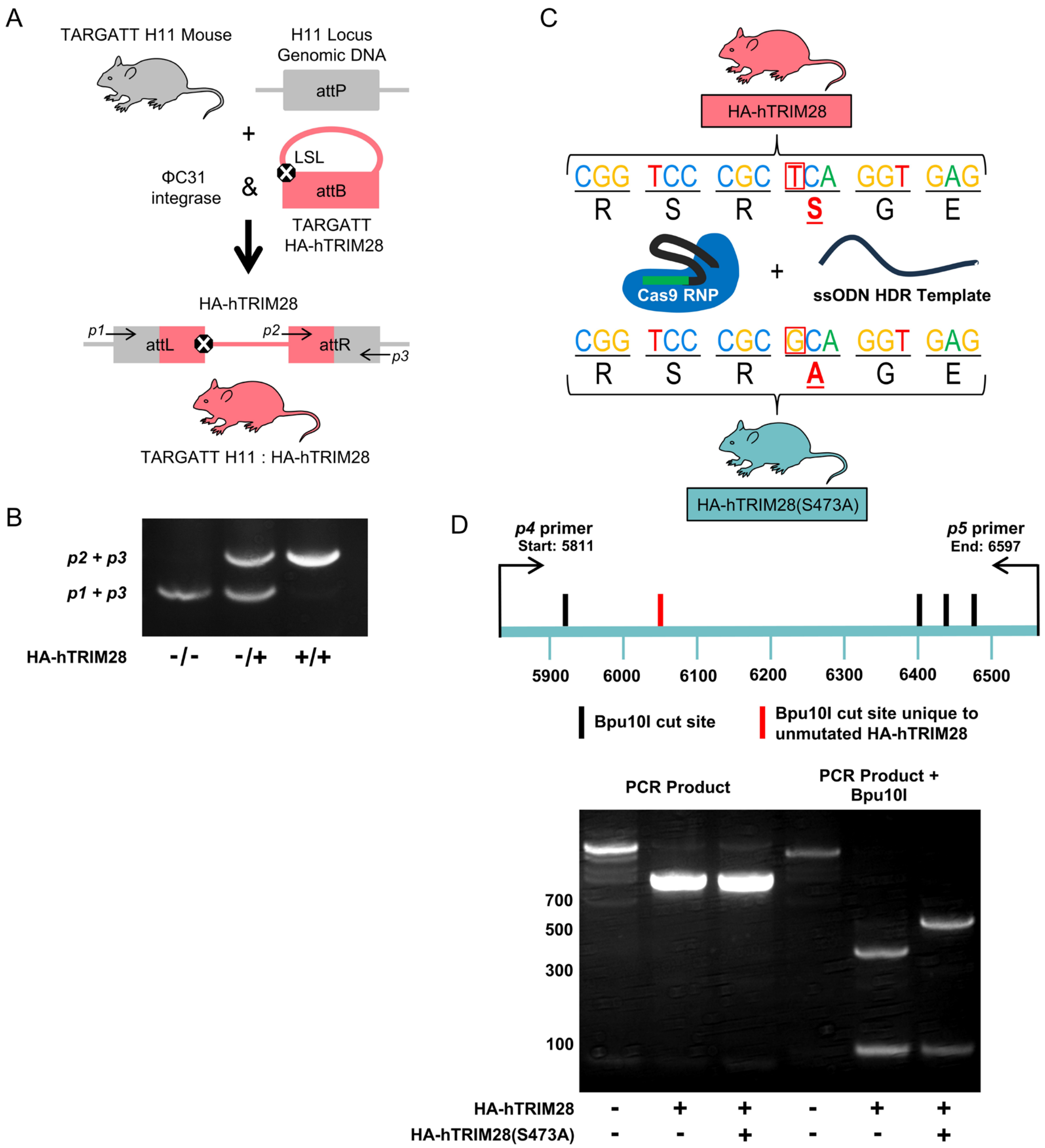


**Fig. S9. Strategy for creating mice that allow for the tamoxifen-inducible expression of human TRIM28 or a S473A phosphodefective mutant of hTRIM28.** (**A**) Embryos from TARGATT mice that contained an attP integration site in the H11 locus were injected with ΦC31 integrase and an HA-hTRIM28 TARGATT vector that contained a LoxP-Stop-LoxP (LSL) cassette and an attB integration sequence. (**B**) As illustrated in A, three primers (*p1*, *p2,* and *p3*) were used to confirm the successful integration of the HA-hTRIM28 vector into the genomic DNA of the offspring. (**C**) CRISPR-Cas9-mediated homologous-directed repair was used to make a single point mutation in TARGATT H11 : HA-hTRIM28 mice that switched the serine 473 residue of hTRIM28 to a non-phosphorylatable alanine (HA-hTRIM28(S473A). (**D**) Primers *p4* and *p5* along with Bpu10I digestion were used to confirm the differences in the genomic DNA of the mice that expressed HA-hTRIM28 versus HA-hTRIM28(S473A).

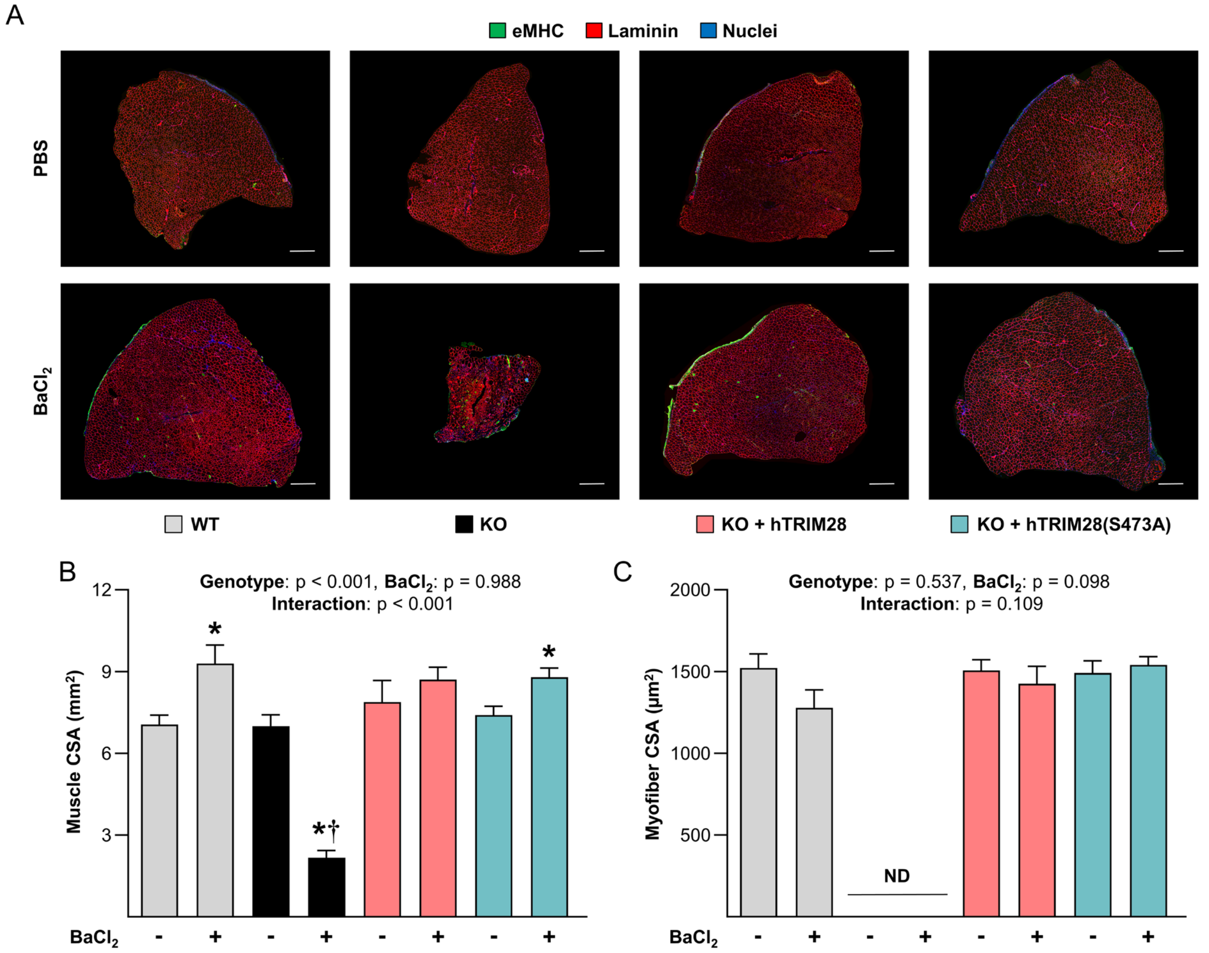


**Fig. S10. TRIM28(S473) phosphorylation in satellite cells is not required for the restoration of muscle CSA or myofiber CSA following $BaCl_2$-induced injury.** At 14 days post tamoxifen the tibialis anterior muscles of wild-type (WT) mice, tamoxifen-inducible and satellite cell-specific TRIM28 knockout mice (KO) mice, as well as KO mice that contain tamoxifen-inducible "rescue" expression of hTRIM28 (KO + hTRIM28) or phosphodefective hTRIM28 (KO + hTRIM28(S473A)) were injected with $BaCl_2$ (+) to induce injury or PBS (-) as a control condition. The tibialis anterior muscles were collected after a 21-day recovery period. (**A**) Mid-belly cross sections of the muscles were subjected to immunohistochemistry for eMHC, laminin, and nuclei. Scale bar = 500 µm. (**B**) Measurements of whole muscle cross-sectional area (CSA), and (**C**) the mean myofiber CSA per muscle. Values are presented as the group mean + SEM, *n* = 6/group. Due to the absence of clear myofibers in the $BaCl_2$-treated muscles of KO mice, myofiber CSA data for these mice was not determined (ND). Data were analyzed with two-way repeated-measures ANOVA. * indicates a significant effect of $BaCl_2$ within a given genotype, † indicates significant difference from the $BaCl_2$ treated WT condition, $p < 0.05$.

| Antibodies | Source | Identifier | Dilution |
|---|---|---|---|
| Anti-Mouse Total Tif1β (Kap-1, TRIM28) (C42G12) | Cell Signaling Technologies (Danvers, MA, USA) | Cat# 4124S | IHC: 1:30<br>WB: 1:1000 |
| Anti-HA-Tag (C26F4) | Cell Signaling Technologies (Danvers, MA, USA) | Cat# 3724S | IHC: 1:300 |
| Anti-Mouse P-Tif1β (KAP-1, TRIM28) Ser473 Poly6446 | BioLegend (San Diego, CA, USA) | Cat# 644602 | IHC: 1:30 |
| Anti-Mouse PAX7 | Developmental Studies Hybridoma Bank (Iowa City, IA, USA) | AB_528428 | IHC: 1:10 |
| Anti-Mouse MYH3 (embryonic myosin heavy chain) (F1.652) | Developmental Studies Hybridoma Bank (Iowa City, IA, USA) | AB_528358 | IHC: 1:350 |
| Anti-Mouse MYH2 (Type IIa myosin heavy chain) (SC-71) | Developmental Studies Hybridoma Bank (Iowa City, IA, USA) | AB_ 2147165 | IHC: 1:100 |
| Anti-Mouse MYH4 (Type IIb myosin heavy chain) (BF-F3) | Developmental Studies Hybridoma Bank (Iowa City, IA, USA) | AB_ 2266724 | IHC: 1:50 |
| Anti-Mouse MYH1 (Type IIx myosin heavy chain) (6H1) | Developmental Studies Hybridoma Bank (Iowa City, IA, USA) | AB_2314830 | IHC: 1:10 |
| Anti-Mouse MHC (all myosin heavy chain isoforms) (MF 20) | Developmental Studies Hybridoma Bank (Iowa City, IA, USA) | AB_2147781 | IHC: 1:50 |
| Anti-Mouse MYOD (5.8A) | BD Pharmingen (Franklin Lakes, NJ, USA) | Cat# 554130 | IHC: 1:50<br>WB: 1:1000 |
| Anti-Mouse Dystrophin (Dy8/6C5) | Novocastra (Leica Biosystems, Buffalo Grove, IL, USA) | Cat# NCL-DYS2 | IHC: 1:100 |
| Anti-Mouse MYOG (F5D) | Santa Cruz (Dallas, TX, USA) | Cat# sc-12732 | IHC: 1:50<br>WB: 1:500 |
| Anti-Mouse Myomixer | ThermoFisher Scientific (Waltham, MA, USA) | Cat# AF4580SP | WB: 1:2000 |
| Anti-Mouse Dystrophin | Abcam (Cambridge, UK) | Cat# ab15277 | IHC: 1:300 |
| Anti-Mouse Myomaker | Abcam (Cambridge, UK) | Cat# ab188300 | WB: 1:200 |
| Anti-Mouse Ki67 (SP6) | Abcam (Cambridge, UK) | Cat# ab16667 | IHC: 1:50 |
| Anti BrdU (BMC9318) | Sigma-Aldrich (St. Louis, MO, USA) | Cat# 11170376001 | IHC: 1:20 |
| Anti-Mouse Laminin | Sigma-Aldrich (St. Louis, MO, USA) | Cat# L9393 | IHC: 1:500 |

| Antibodies | Source | Identifier | Dilution |
|---|---|---|---|
| Peroxidase-labeled Anti-Sheep Secondary | Sigma-Aldrich (St. Louis, MO, USA) | Cat# A3415 | WB: 1:1000 |
| Fab Fragment Goat Anti-Mouse IgG (H+L) Block | Jackson Immunoresearch (West Grove, PA, USA) | Cat# 115-007-003 | IHC: 1:10 |
| AMCA Anti-Mouse IgM | Jackson Immunoresearch (West Grove, PA, USA) | Cat# 115-155-075 | IHC: 1:150 |
| Alexa Fluor 488 Anti-Mouse IgG1 | Jackson Immunoresearch (West Grove, PA, USA) | Cat# 115-545-205 | IHC: 1:2000 |
| FITC Anti-Mouse IgG2b | Jackson Immunoresearch (West Grove, PA, USA) | Cat# 115-095-207 | IHC: 1:200 |
| Alexa Fluor 594 Anti-Mouse IgG1 | Jackson Immunoresearch (West Grove, PA, USA) | Cat# 115-585-205 | IHC: 1:2000 |
| Alexa Fluor 488 Anti-Rabbit IgG | Invitrogen (Carlsbad, CA, USA) | Cat# A11008 | IHC: 1:5000 |
| Alexa Fluor 594 Anti-Rabbit IgG | Invitrogen (Carlsbad, CA, USA) | Cat# A11037 | IHC: 1:5000 |
| Peroxidase-labeled Anti-Rabbit Secondary | Vector Labs (Burlingame, CA USA). | Cat# PI-1000 | WB: 1:5000 |
| Peroxidase-labeled Anti-Mouse Secondary | Vector Labs (Burlingame, CA USA). | Cat# PI-2000 | WB: 1:5000 |

**Table S1. List of antibodies used for immunohistochemical (IHC) and western blot (WB) analyses**
